# A new targeting motif for endoplasmic reticulum surface proteins

**DOI:** 10.1101/2024.04.22.590521

**Authors:** Sivan Arad, Parkesh Suseendran, Shani Ravid, Yoav Peleg, Shifra Ben-Dor, Elena Fidel, Tsviya Olender, Xiaofeng Wang, Maya Schuldiner, Emma J Fenech

## Abstract

The Endoplasmic Reticulum (ER) is the entry site to the secretory pathway, serving as the targeting destination for ∼30% of the proteome. The mechanisms for targeting soluble or integral membrane secretory pathway proteins are well-studied. However, it is currently unknown how the tens of ER surface proteins (SuPs), central for organelle function, reach the outer leaflet of the membrane. It was previously shown that an amphipathic helix (AH) from the Brome mosaic virus protein 1a, is both necessary and sufficient for targeting to the ER surface in baker’s yeast. We therefore utilized this helix as a model substrate and performed a high-content screen to uncover factors that affect SuP targeting. Our results suggest a role for membrane lipid composition in targeting specificity. To see if the presence of an AH is a more general mechanism for SuP targeting, we searched for their presence within SuPs of both yeast and humans. Five endogenous yeast SuPs contained AHs, and of these four were sufficient for ER localization. Moreover, the presence of an AH was conserved to human SuP orthologs. By altering helix features we determine the parameters that affect this new targeting motif. Hence our work demonstrates how specific properties of AHs encode affinity for the ER membrane. More globally, understanding how SuPs are targeted correctly takes us a step forward in determining the underlying mechanisms of cellular localization and secretory pathway functions.

The authors declare that they have no conflict of interest.

## Introduction

The Endoplasmic Reticulum (ER) is a multifunctional and essential organelle. Crucially, it serves as the entry site and maturation environment for both membranal and soluble secretory pathway proteins. In the baker’s yeast (*Saccharomyces cerevisiae*), a third of the proteome (around 2000 proteins (Chen et al., 2005; Choi et al., 2010; Wallin and Heijne, 1998)) is targeted to the ER, where they either remain as residents or are sorted to downstream secretory pathway organelles. All these proteins must target correctly to the ER to support a functional secretory pathway that meets cellular demands.

A lot is known about the ER targeting mechanisms of soluble and membranal secretory pathway proteins. Such proteins can be targeted by three highly conserved pathways: the SRP (Signal Recognition Particle) (Walter et al., 1981; Walter and Blobel, 1981a, 1981b); GET (Guided Entry of Tail-anchored proteins) (Schuldiner et al., 2008; Stefanovic and Hegde, 2007); or SND (SRP-independent) (Aviram et al., 2016) pathways. Every targeting pathway has a unique cargo range that depends on the presence of different targeting motifs (signal peptides or transmembrane domains (TMDs)) and their exact location within the protein (the amino (N’) or carboxy (C’) terminus or in a central position (Aviram and Schuldiner, 2017)). Regardless of these parameters, to date, all characterized pathways target proteins that have a hydrophobic domain that enables either translocation into the lumen (in the case of the signal sequence) or insertion into the membrane (in the case of a TMD). However, there are proteins that must be targeted to the ER membrane but that do not translocate into its membrane or lumen – ER surface proteins (SuPs). These proteins do not contain a signal sequence or TMDs, yet they are somehow specifically targeted to the outer leaflet of the ER membrane where they function.

In this work we set out to uncover what guides the targeting of the tens of ER SuPs, many of which are crucial for secretory pathway function. We initially focused on a well-characterized ER SuP that is encoded by the Brome mosaic virus (BMV), a positive-strand RNA virus that primarily infects monocotyledon plants (Wang and Ahlquist, 2008). BMV replication can be recapitulated in baker’s yeast (Ishikawa et al., 1997) where it forms viral replication complexes at the perinuclear ER membrane to multiply itself. During this process, BMV replication protein 1a recruits host proteins involved in lipid synthesis/modification or membrane shaping to invaginate the outer ER membrane into the lumen. BMV 1a additionally recruits viral replicase 2a^pol^ and viral genomic RNAs as replication templates (Liu et al., 2009; Wang and Ahlquist, 2008; Diaz and Wang, 2014; He et al., 2021). To do all this 1a must reach the ER membrane yet it does not harbor a TMD. Rather, 1a contains two amphipathic helices (AHs), Helix A and B, that are important for its localization. Moreover, Helix B (1aHB) has been shown to be both necessary and sufficient to target to the ER surface (Sathanantham et al., 2022).

An AH is a helix with two faces – hydrophobic and hydrophilic, that are kept constant across the helix axis. This property enables the helix to interact with membrane phospholipids on the one side, while interacting with the cytosolic milieu on the other. AHs similar to 1aHB are found among more viruses from the *Alsuviricetes* class (Ahola and Karlin, 2015), such as cowpea chlorotic mottle virus (CCMV), cucumber mosaic virus (CMV), hepatitis E virus (HEV), and Rubella virus (RuV) (Sathanantham et al., 2022). These viral-derived AHs target fluorescence proteins to specific organelles where these viruses assemble their viral replication complexes (Sathanantham et al., 2022). Moreover, AHs in cellular proteins have been shown to enable specific targeting to other organelles such as lipid droplets (Čopič et al., 2018) and the Golgi apparatus (Magdeleine et al., 2016; Bigay et al., 2005). Taken together these data suggest that AHs can indeed function as a specificity determinant for targeting.

We first performed a genetic screen in yeast to find mutations that affect localization of 1aHB. The results of this screen suggested that the ER lipid composition is crucial for the helix to localize specifically to the ER membrane. Then, we searched for AHs within endogenous yeast and human SuPs, and showed many such AHs exist, some conserved in their presence from yeast to human. Importantly, the endogenous yeast AHs were also sufficient for specific ER targeting. To better understand how these AHs target specifically to the ER, we examined several characteristics and found both helix length and the presence of phosphorylatable residues to be important. Altogether we describe a new targeting motif for endogenous SuPs and provide an initial glimpse into the elements that regulate SuP targeting.

## Results

### A high-content screen suggests that ER surface targeting is regulated by lipid composition

We set out to uncover how SuPs, which are localized to and function on the outer ER leaflet, are targeted (Figure 1A). To discover factors in trans that affect SuP ER targeting, we used the well-characterized AH, 1aHB (Figure 1B), as a model. When fused to Green Fluorescent Protein (1aHB-GFP) we could confirm that this AH is sufficient to confer ER targeting since it colocalized with an ER marker (Figure 1C) (Sathanantham et al., 2022). We then performed a genetic screen and looked for genes whose mutation altered the localization of 1aHB. Genomically integrated 1aHB-GFP was introduced into mutant collections (libraries) where each strain harbors a deletion of one non-essential gene (Giaever et al., 2002) or a hypomorphic allele of an essential one (Decreased Abundance by mRNA Perturbation, DAmP) (Breslow et al., 2008). We screened these libraries using fluorescence microscopy, analyzed the images, and identified candidate genes that affect localization – defined as strains with a different 1aHB-GFP phenotype relative to the control strain (Figure 1D).

**Figure 1.**
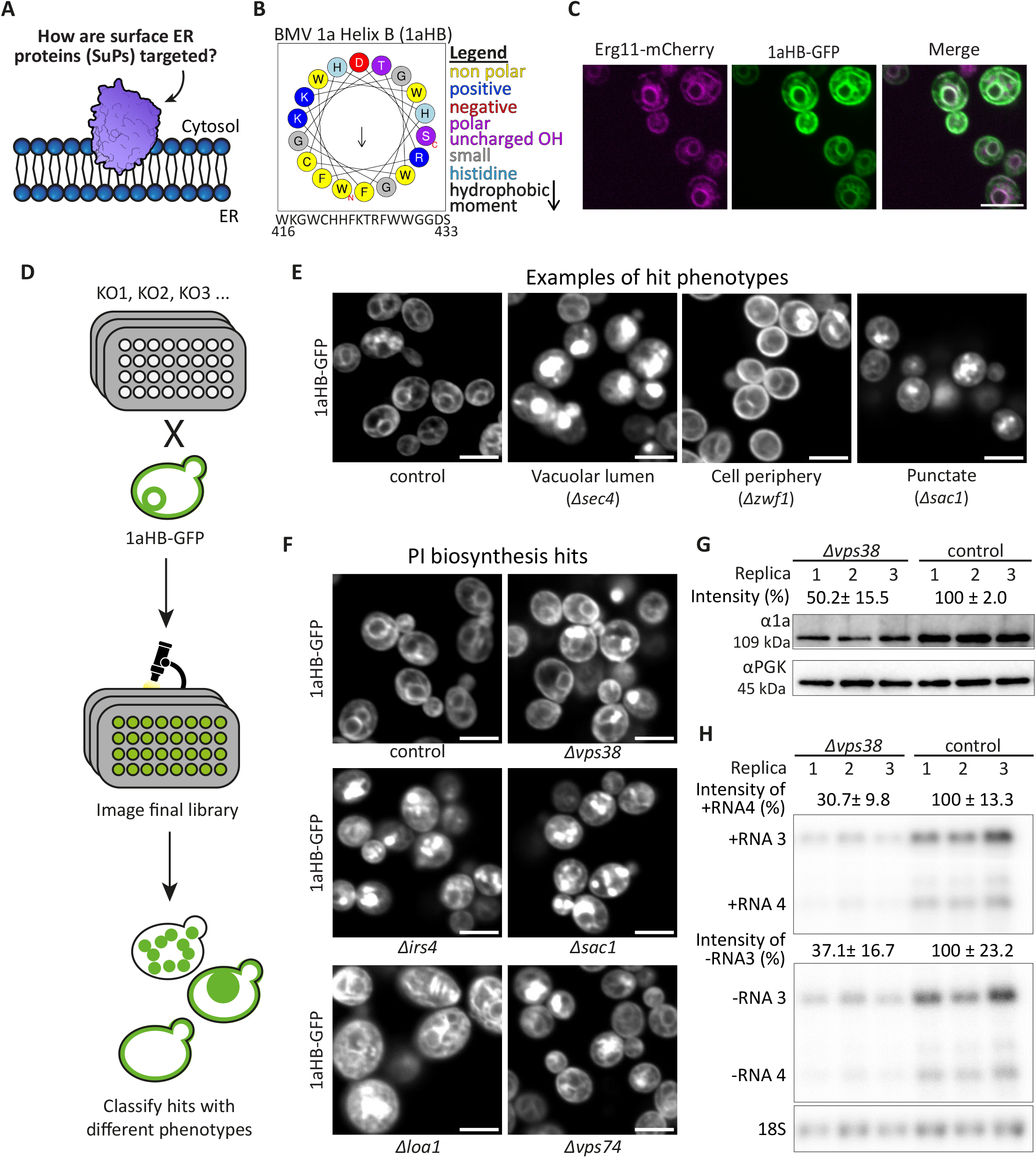
A high-content screen suggests that ER surface targeting is regulated by lipid composition. (**A**) A schematic diagram of the research question: How are ER SuPs targeted correctly? (**B**) A helical wheel of BMV 1a Helix B (1aHB) shows that this helix is amphipathic. A legend of the amino acid groups is included. The hydrophobic moment value is presented in Table S5. (**C**) Fluorescent microscopy images of a strain expressing the ER marker Erg11-mCherry and 1aHB fused at its C’ to GFP. The images show a typical ER localization pattern and colocalization, confirming that the helix is sufficient for encoding targeting information. (**D**) A schematic diagram of the screening process. A systematic high-content microscopy screen was performed after integrating 1aHB-GFP into the background of a whole-genome library of deleted (Giaever et al., 2002) or decreased-expression (Breslow et al., 2008) proteins. The resulting collection was imaged and mutants where 1aHB was mislocalized were identified. (**E**) Fluorescent microscopy images representing the three types of hit phenotypes observed. From left to right: (1) increased vacuolar lumen accumulation, (2) appearance of a signal at the cell periphery, and (3) accumulation of punctate structures. In parentheses are the actual representative strains from the original screen. (**F**) Fluorescent microscopy images from the screen of mutant strains that were deleted for genes whose protein product plays a role in phosphoinositide (PI) biosynthesis and were identified as hits. (**G**) Western blot analysis of the levels of protein 1a in BMV-infected yeast strains. A *Δvps38* strain was compared to a control strain and found to have half the levels of 1a protein. PGK levels were assayed as housekeeping protein control, to ensure equal loading. The average values are calculated from four biological replicates, each containing three technical repetitions. (**H**) A BMV replication assay – a Northern blot analysis of the RNA levels of -RNA3 and +RNA4 strands. A deletion of *VPS38* was compared to a control strain. 18S levels were assayed as housekeeping RNA control, to ensure equal loading. The intensity of the tested RNAs in *Δvps38* strain are reduced to 30.7% and 37.1% for +RNA4 and -RNA3, respectively, compared to the control strain. This demonstrates a reduced capacity of BMV to replicate in strains with reduced PI3P. The average values are calculated from four biological replicates, each containing three technical repetitions. A representative replicate is presented in the figure. The scale bar in all the microscopy images is 5µm.

We found 136 mutant strains that had different subcellular distribution patterns compared to the control (Table S1) (Figure 1E). Interestingly, we did not find any hits that represent components of the known ER targeting machineries (such as GET, SRP or SND genes). While there are many reasons to miss phenotypes in a systematic screen, these findings suggest that an additional mechanism may be at play. This notion is supported by the fact that no mutant strain caused 1aHB to become completely mislocalized to the cytosol, but rather 1aHB was displaced to non-ER membranes or aggregates (Figure 1E).

We noticed that 19 out of the 136 mutants are proteins involved in phosphatidylinositol (PI) or other lipid biosynthesis processes (Figure 1F). PI is a major phospholipid in membranes and could directly influence the properties of the bilayer. To further examine whether mutants in PI biosynthesis really affect 1a localization we tested the capacity for BMV to replicate in the absence of *VPS38. VPS38* is part of the Vps34 PI3-kinase complex, and affects ER PI3P levels (Kihara et al., 2001). Indeed, we found that in a *Δvps38* strain, the relative intensity of 1a is reduced by half (Figure 1G). Moreover, viral replication was reduced to a third based on the accumulation of negative strand (-)RNA3 (a replication intermediate) and positive strand (+)RNA4 (a product of BMV viral replication) (Figure 1H).

The screen results, together with the viral-based assays, suggest that lipid composition affects the HB-mediated 1a targeting to the ER. The absence of PI or its phosphorylated derivatives such as PI3P, could decrease 1a’s affinity to the ER, or increase its affinity to a non-ER membrane,thus reducing its overall levels and resulting in decreased viral capacity to replicate and infect other cells.

### Predicting AHs in endogenous yeast proteins uncovers a new ER-targeting motif

The 1aHB screen (Figure 1) suggested that the direct interaction between the ER membrane and a viral AH is crucial for correct targeting. We therefore wanted to understand if native AHs could also play a role in ER targeting of endogenous yeast SuPs. To do this we first created a list of all yeast proteins that are ER localized (Yofe et al., 2016; Weill et al., 2018; Huh et al., 2003) yet do not have either a known or predicted signal sequence, or a TMD (Weill et al., 2019), and hence are likely to remain as SuPs on the cytosolic face of the ER membrane. This analysis resulted in 49 putative SuPs (See Table S2). Not surprisingly, some of these proteins are soluble members of large complexes that span the ER membrane and therefore may correctly localize through interactions with their complex components. Examples of this include the known soluble targeting factors of the GET and SRP pathways, e.g., Get3 and Srp54. However, for the majority of the SuPs, the targeting mechanism was unknown. Hence, we used Heliquest (Gautier et al., 2008) (see Methods) to predict the presence of AHs in these proteins (Figure 2A).

**Figure 2.**
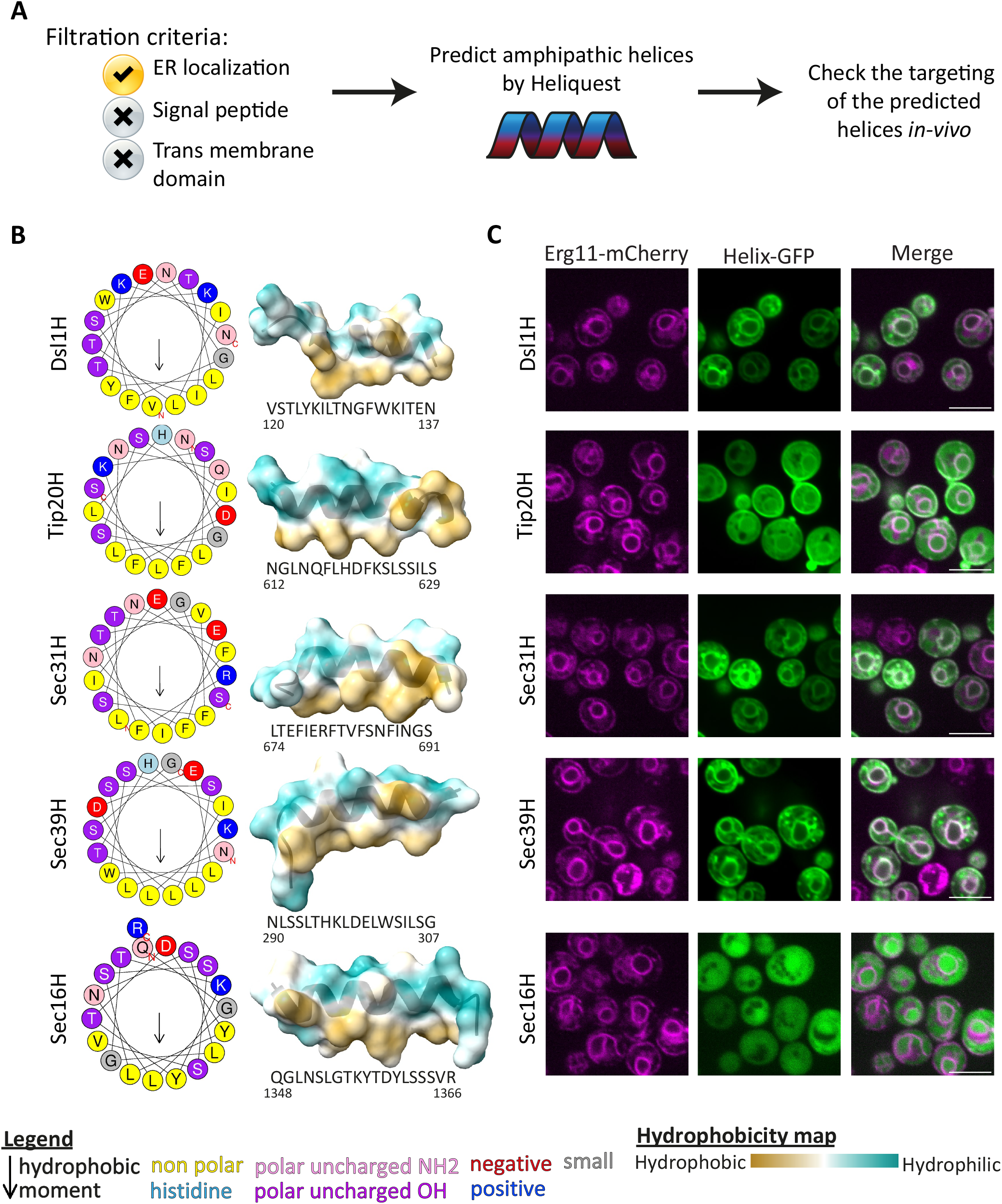
Predicting AHs in endogenous yeast proteins uncovers a new ER-targeting motif. (**A**) A schematic diagram of the process that was performed to find new ER targeting domains. Proteins that are localized to the ER when fused to a GFP tag (Yofe et al., 2016; Weill et al., 2018; Huh et al., 2003), but have no signal sequence or TMD (Weill et al., 2019), were shortlisted, given that they have no defined ER targeting pathway. Heliquest (Gautier et al., 2008) was then used to predict the presence of an AH such as HB of the 1a BMV protein. Expressing the helices fused to GFP uncovers their ER surface targeting capacity. (**B**) Helical wheels and 3D modeling of the predicted helices, which are all amphipathic. Hydrophobic moment values are presented in Table S5. (**C**) Fluorescent microscopy images of strains that express the ER marker Erg11-mCherry and the predicted helices fused to GFP at their C’. The images show ER localization for all helices except Sec16H, suggesting that they are sufficient for targeting. The scale bar is 5µm.

From all ER SuPs, we predicted AHs in Dsl1, Tip20, Sec31, Sec39, and Sec16 (which will be referred to as “(protein name)H” (Figure 2B). To determine whether these helices are sufficient for ER targeting, we generated plasmids encoding the helices fused at their C’ with GFP. These plasmids were then introduced into a strain genomically encoding a red ER marker (Erg11-mCherry) to assess co-localization (Figure 2C). To better recognize subtle differences in localization, we used enhanced resolution imaging (see Methods). From our results, we could see that most of the AHs are indeed localized to the ER, either fully or partially. The only AH that was exclusively cytosolic and nucleoplasmic was Sec16H. A possible explanation for this is that the hydrophobic face of this AH is not fully continuous (Figure 2B), unlike the other AHs that have a longer continuous hydrophobic patch.

Put together, our analysis uncovered four endogenous ER proteins that have an AH that is sufficient to confer localization to the ER membrane, suggesting that they play a role in the targeting of these proteins.

### ER targeting of AHs relies on their biochemical properties

The hits that arose in the 1aHB screen (Figure 1F), suggest that ER targeting of the helices is governed by their biochemical properties which would confer an affinity between the helix and the ER membrane lipids. Since the endogenous AHs are also sufficient for ER targeting and they share many biochemical traits with 1aHB (Figures 1 and 2), we hypothesized that they may share conserved targeting elements.

If the affinity to the ER membrane is indeed based on the biochemical properties of the sequence, then the precise order of amino acids should not be important, and we would expect it to be less conserved. However, we found that the features (hydrophobicity, charge, etc.) of the first nine amino acids (aa) of Sec31H are strongly conserved in related neighboring fungi (Figure 3A, Table S3). To examine if this part of the Sec31H sequence is a sufficient targeting motif, we visualized a GFP tagged version of this domain but found that it remains cytosolic, suggesting that a simple protein motif is not enough to explain the targeting (Figure 3B).

**Figure 3.**
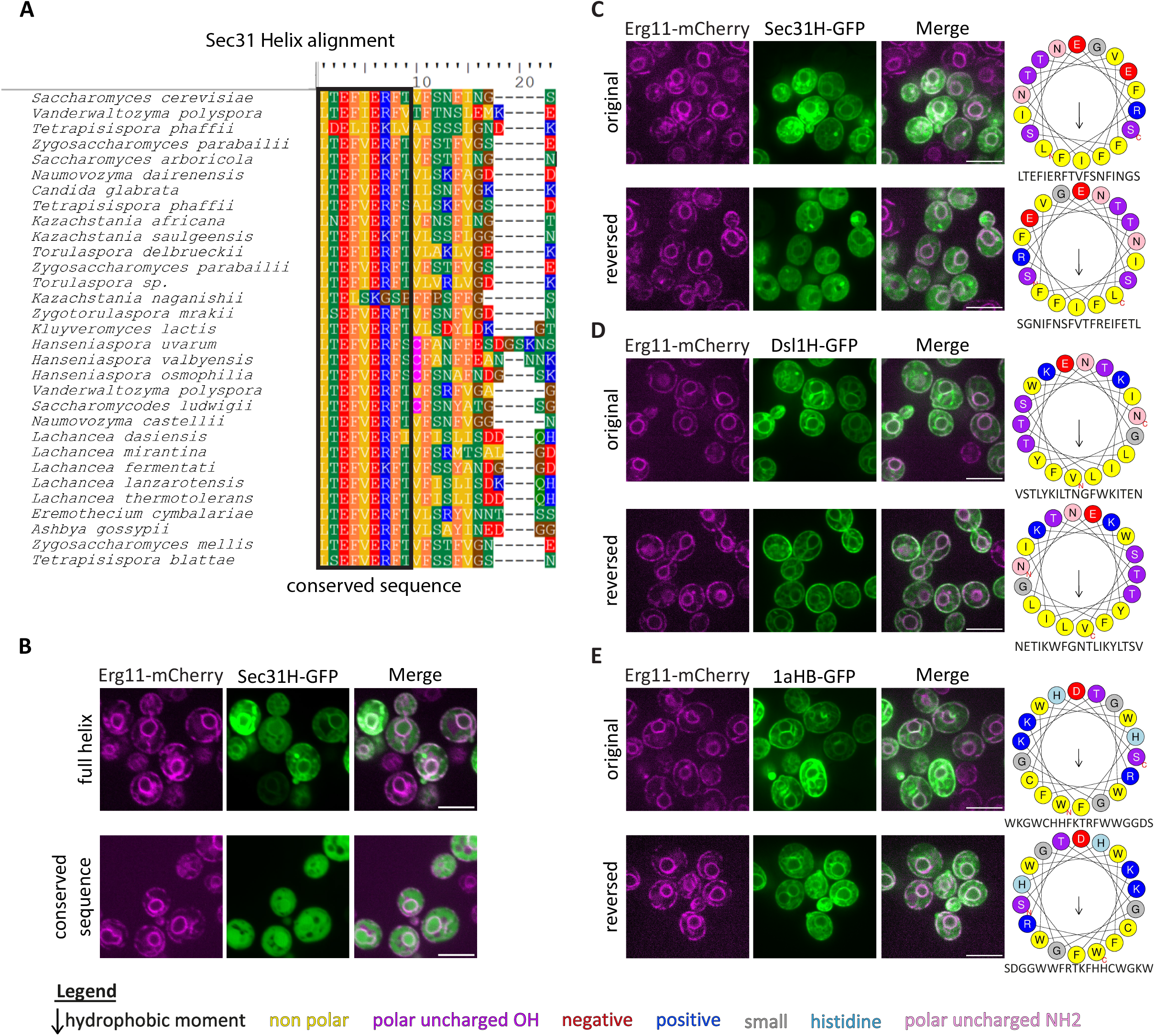
ER targeting of AHs relies on their biochemical properties. (**A**) The conservation of Sec31H domains across similar species. The first nine amino acids are highly conserved. (**B**) Fluorescent microscopy images of yeast expressing either the full-length or only the conserved region of Sec31H fused to GFP at their C’, and the ER marker Erg11-mCherry. The highly conserved region is not sufficient for ER targeting. (**C**) – (**E**) Fluorescent microscopy images of yeast expressing the ER marker Erg11-mCherry and various helices fused with GFP at their C’. To the side of each microscopy panel, a helical wheel of the imaged helix is depicted. Hydrophobic moment values are presented in Table S7. The mutated helices have the same amino acids as their native counterparts, spaced similarly, but in a reversed order. The helices are (**C**) Sec31H, (**D**) Dsl1H, and (**E**) 1aHB. The localization of the mutants was compared to the native ones. No major effects on targeting were observed when reversing the amino acids order. The scale bar in all the microscopy images is 5µm.

To further examine if the specific sequence of the whole helix or its biochemical properties are important for ER targeting, we designed helices with a reversed amino acid (aa) order, as previously used (Pranke et al., 2011). In this way, the biochemical properties of the helix are preserved, but helical targeting that is based on an aa motif would be destroyed. We tested three helices with a strong ER affinity and found that in all of them targeting to the ER was maintained (Figure 3C-E).

This data, together with the screen observation that the ER membrane composition is crucial for correct targeting (Figure 1D-F), suggest that indeed, the biochemical properties of the helix are a major determinant for ER targeting and that this may define the affinity to the unique ER lipid composition.

### AH length is a determining factor of its cellular localization

Since AHs have been observed as targeting determinants for other organelles (Magdeleine et al., 2016; Bigay et al., 2005; Čopič et al., 2018), it remained a mystery as to how an AH, that is inherently drawn to various hydrophobic membranes, can govern specificity. To understand which features of our AHs are responsible for defining ER specificity, we turned to analyze and test the properties of this domain that are shared between the different helices; both 1aHB and the endogenous helices (Figures 1B and 2B). One property is the helix length – all of them are exactly 18 aa long. It has previously been shown that longer AHs (25-38 aa length) are targeted to different organelles, such as the Golgi (Magdeleine et al., 2016; Bigay et al., 2005) and lipid droplets (Čopič et al., 2018).

To test the effect of helix length on cellular localization, we designed helices that would maintain the biochemical features of the original ones but be doubled in length. 18 aa form a complete helical wheel, so to preserve the hydrophobic and hydrophilic faces, we duplicated the sequences to make 36 aa long helices, similar to those that confer targeting to other membranes. Remarkably, despite having identical biochemical properties, the GFP tagged double-length helices dramatically changed localization (Figure 4). The long versions of Dsl1H, Sec31H and 1aHB now had only a minor fraction localized to the ER (Figure 4 A, B, D). For the doubled Dsl1H and Sec31H, they both appeared to form punctate structures, with the ones from Dsl1H occupying a large proportion of the cytosol (Figure 4 A, B). Interestingly, the doubled Tip20H and 1aHB appeared much more peripheral relative to their original-length counterparts (Figure 4 C, D), with the latter seeming exclusively peripheral (Figure 4C).

**Figure 4.**
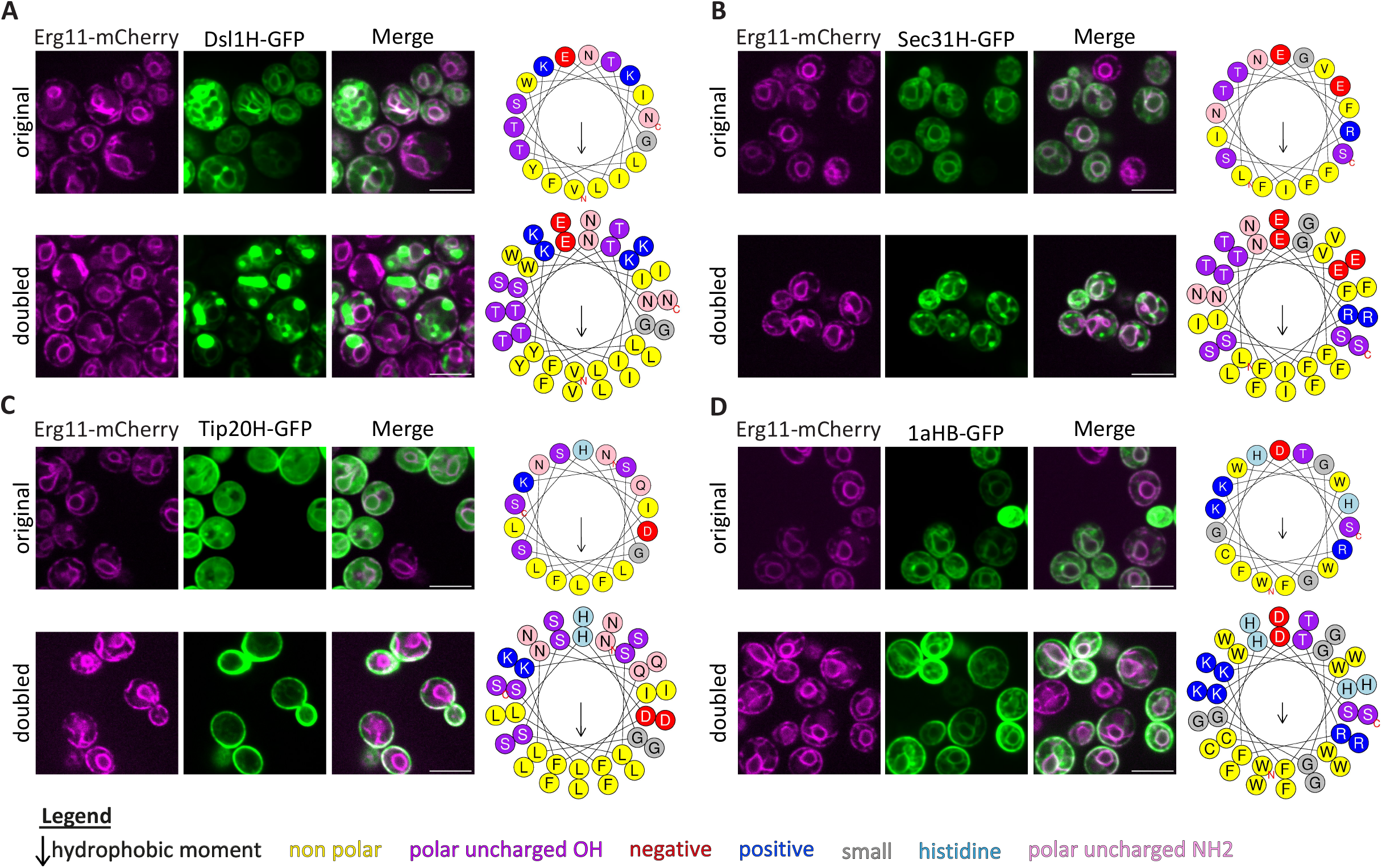
AH length is a determining factor of cellular localization. Fluorescent microscopy images of yeast expressing original- or double-length helices fused to GFP at their C’, and the ER marker Erg11-mCherry. To the side of each microscopy panel, a helical wheel of the imaged helix is depicted. Hydrophobic moment values are presented in Table S5. The mutated helices are twice as long as their native counterparts. The helices are (**A**) Dsl1H, (**B**) Sec31H, (**C**) Tip20H, and (**D**) 1aHB. The localization of the mutants was compared to native, 18aa, ones. The increased length of the helix reduced the ER localization of all helices to various extents. The scale bar in all the microscopy images is 5µm.

These findings demonstrate how important helix length is for determining membrane specificity. Further work is required to examine the new locations of the long helices, and to understand why localization shifts occur to two different locations.

### Bulky residues in the AHs do not define membrane specificity

From analyzing the residues within the amphipathic helices, we found that all the hydrophobic faces have at least one, and usually more, aromatic amino acids such as phenylalanine (Phe), tyrosine (Tyr) or tryptophan (Trp) (Figures 1B and 2B). These residues make these faces bulkier, and this feature may play a role in specifying targeting to the ER, potentially by enabling interactions with the less densely packed lipid composition of the ER membrane (van Meer et al., 2008; Bigay and Antonny, 2012). To test this hypothesis, we mutated the helices to swap all the Phe, Tyr and Trp residues with the compact, hydrophobic aa leucine (Leu), thus conserving hydrophobicity but reducing the bulkiness. When visualizing the GFP tagged mutated helices, we could not find any effect on their ER localization (Figure 5A-C).

**Figure 5.**
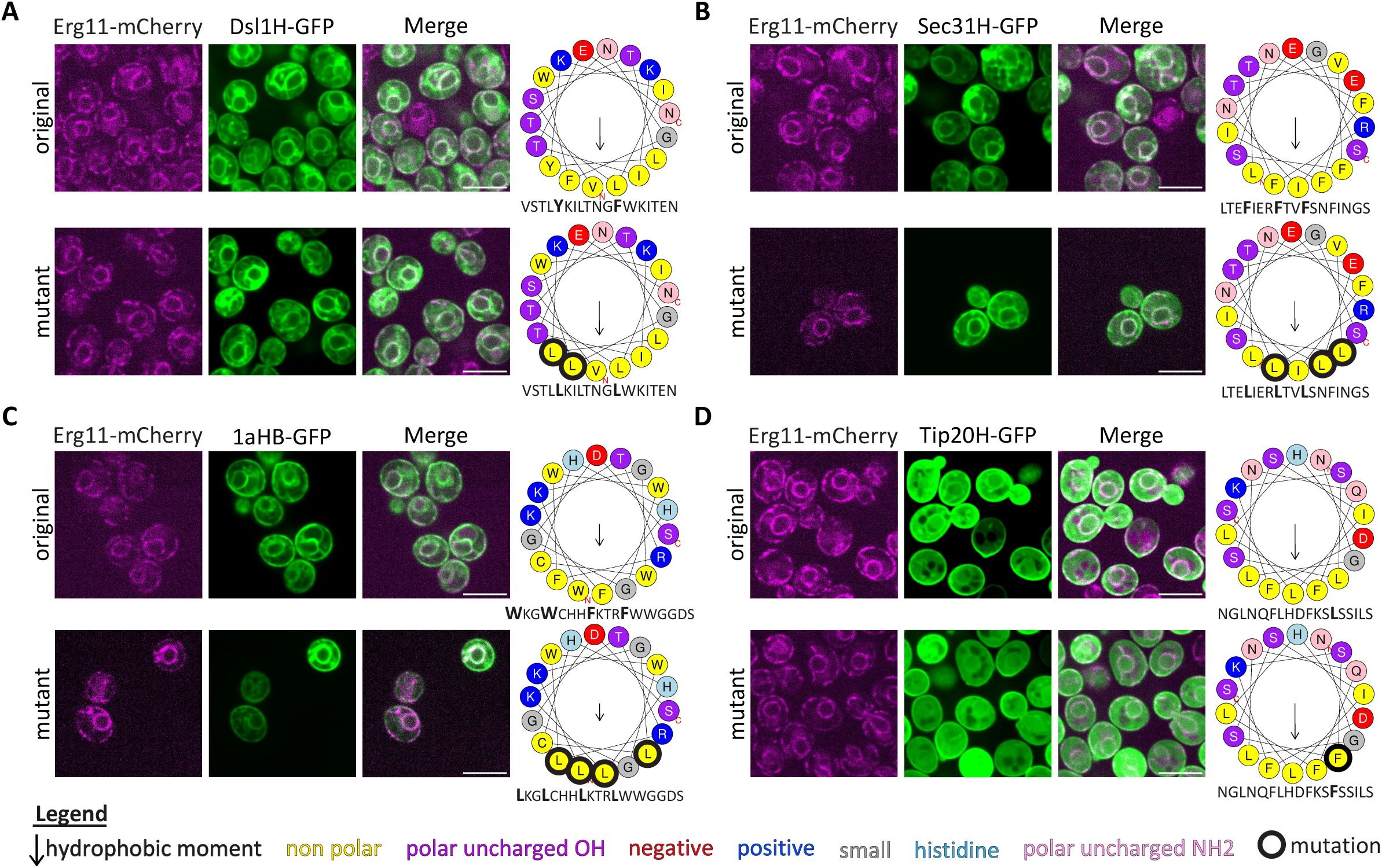
Bulky residues in the AHs do not define membrane specificity. Fluorescent microscopy images of yeast expressing the ER marker Erg11-mCherry and helices with an altered number of bulky residues, fused to GFP at their C’. To the side of each microscopy panel, a helical wheel of the imaged helix is depicted. Hydrophobic moment values are presented in Table S5. The outline of the mutations is marked in bold. In panels (**A**)-(**C**), the aromatic residues were replaced with small hydrophobic residues and the helices are (**A**) Dsl1H, Sec31H, (**B**) and (**C**) 1aHB. In panel (**D**), the opposite mutation on Tip20H is presented, where one small hydrophobic residue was replaced with an aromatic residue. The localization of the mutants was compared to their controls. No major effects on targeting were observed by removing or adding aromatic residues. The scale bar in all the microscopy images is 5µm.

The bulkiness reduction showed that this feature may not be necessary for ER specificity, but maybe it acts as a contributing property that, to some extent, can be sufficient for ER targeting. To test this idea, we focused on Tip20H, which has the weakest ER affinity (Figure 2C, second row) and was hence not included in the above perturbations. We then mutated it in the inverse manner, to increase the number of bulky residues, by replacing Leu with Phe at the 13^th^ position (Figure 5D). Interestingly, this mutant of Tip20H presented an even more cytosolic localization in comparison to its original sequences (Figure 5D).

Overall, these results suggest that while bulkiness of the hydrophobic face of the helix may play a subtle role, it is unlikely to be the main effector for ER surface specificity of the AHs.

### AH charge may regulate targeting

An additional shared property of the helices is that all of them have several serine (Ser) and threonine (Thr) residues, or negatively charged residues, next to the hydrophobic patch or within the adjacent hydrophilic surface (Figures 1B and 2B). Both Ser and Thr residues can either be phosphorylated to become negatively charged or be in a non-phosphorylated, neutral, state. Previous work has shown that replacing Ser/Thr at the ends of the hydrophobic patch of BMV 1a helix A (not the HB that we focus on here), resulted in mislocalization to the cytosol in yeast cells (Liu et al., 2009) supporting their importance for AH localization.

To test the contribution of these residues to the AH specificity, we started by replacing all the Ser and Thr residues with similar ones that cannot be phosphorylated: alanine (Ala) and valine (Val), respectively. Imaging the GFP tagged mutant helices we found no significant effect on Dsl1H and 1aHB (Figure 6A-B). However, for both Sec31H and Tip20H, the localization was shifted from the ER to the cell periphery, with a few internal puncta also observed (Figure 6C-D, first and third rows).

**Figure 6.**
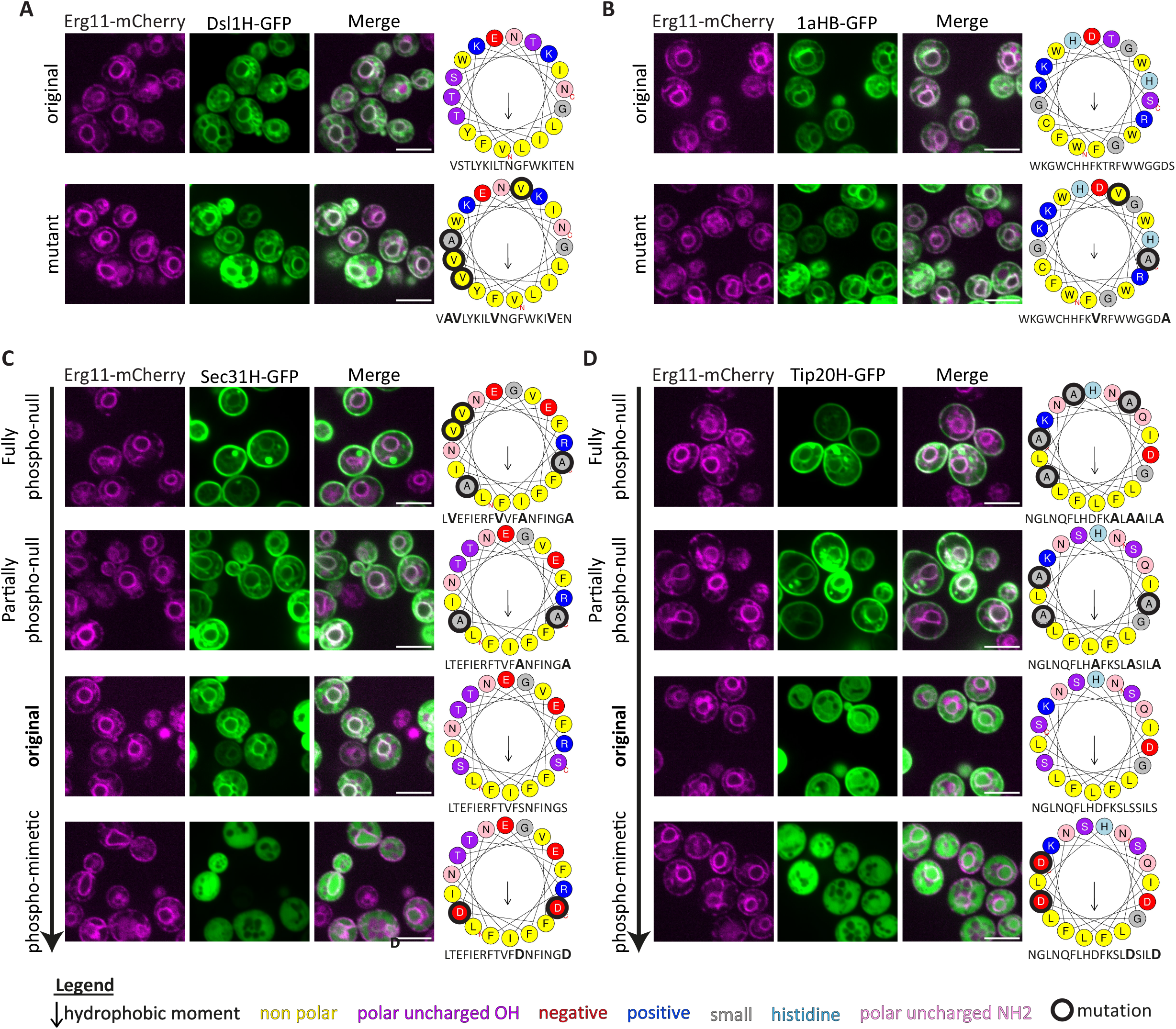
AH charge may regulate targeting. Fluorescent microscopy images of yeast expressing the ER marker Erg11-mCherry and helices with different phosphorylation properties, fused to GFP at their C’. A helical wheel of the imaged helix is shown on the side of each microscopy panel. Hydrophobic moment values are presented in Table S5. The outline of the mutations is marked in bold. (**A**)-(**B**) The mutated helices of Dsl1H and 1aHB, respectively, have phosphor-null mutations of Ser to Ala and Thr to Val. The localization of the mutants was compared to their controls and no major effects on targeting were observed. (**C**)-(**D**) A series of mutations of Sec31H and Tip20H were tested, making a gradient from phosphor-null (rows 1-2 in each panel) to having a constant charge (phosphor-mimetic, row 4 in each panel) around their hydrophobic face. The phenotypes show a gradient of localizations from the cell periphery and the ER to the cytosol. The scale bar in all the microscopy images is 5µm.

Since both Sec31 and Tip20 were sensitive to changes in Ser/Thr composition, we further examined if a charge ‘gradient’ may explain the difference in the cellular localization. For both helices, we tested two more mutant classes. The first class is a partial phospho-null mutant, in which the residues proximal to the hydrophobic patch are unable to be phosphorylated (and therefore are always neutral), by replacing Ser and aspartic acid (Asp) residues to Ala (Figure 6C-D, second rows). The second class is a phospho-mimetic mutation, in which the Ser residues in proximity to the hydrophobic patch were replaced with Asp, to test the effect of having a constant negative charge at these residues in the helix (Figure 6C-D, bottom rows). Indeed, these helix mutants form a gradient of localizations, ranging from peripheral (complete phospho-null mutants), to more ER (partially phospho-null mutant + WT), and to the cytosol (phospho-mimetic mutants) (Figure 6C-D). This observation suggests that charge affects helix localization, and that the phosphorylation state may play an important role in regulating AH targeting to the ER.

## Discussion

In this work, we shed light on the targeting parameters for a subgroup of ER SuPs. We predicted five endogenous AHs in yeast ER SuPs that have similar properties to 1aHB, and four of them were verified *in-vivo* to be sufficient for targeting to the ER. All of these proteins play a role in, or are associated with, COPI (Kraynack et al., 2005) and COPII (Salama et al., 1997) trafficking that takes place between the ER and the Golgi. Understanding their targeting strategies can therefore shed light on inter-organelle interactions.

The presence of AHs in some of the endogenous proteins that we found was surprising at first. Both Dsl1 and Sec31 are well-studied, their structures in a complex have been solved and clearly show that they are not in direct contact with the ER membrane. Rather, Dsl1 and Sec31 interact with their complex partners: Tip20 and Sec13, respectively (Sec31 can also interact with Sec23/24) (Tripathi et al., 2009; Ren et al., 2009; Hutchings et al., 2021). If these proteins do not bind the ER in their final conformation, why would they need an ER targeting motif? We suggest that these ER SuPs have a primary targeting step, that supports the protein to reach the correct membrane. This primary targeting then increases the chances to find the correct binding partner and is then followed by a separate, maintenance stage – where the protein is held in functional complexes at its destination.

Indeed, according to structural data of the Sec31-Sec13 complex (PDB entry number: 2PMS), the AH is buried inside the structure of Sec31 when it is in complex with Sec13 (Fath et al., 2007). We therefore hypothesize that immediately following translation, the AH is exposed since Sec31 is not yet folded. It is at this stage that the AH would serve as the ER targeting component. Then, once the protein arrives at its destination membrane and interacts with its heterodimeric complex member, Sec13, the targeting domain is not required and a conformational change would occur, burying the AH inside the protein core. Further experiments are clearly required to confirm this theory. Remarkably however, AH predictions for human ER SuPs, suggest that about 19 of them include an AH (Table S4). In particular, the human homologs of Sec31 and Dsl1, SEC31B and ZW10, respectively, also contain predicted AHs (Figure S1). The conservation of the presence of AH domains in ER SuPs, despite the helix sequences themselves not being conserved, supports their importance in the biogenesis of these proteins.

It is hard to imagine how a simple AH could confer specificity to a singular membrane. However, previous work on AHs has shown that they are specific to certain membranes such as Golgi cisternae (Magdeleine et al., 2016; Bigay et al., 2005) and lipid droplets (Čopič et al., 2018). Moreover, the AHs that we found all target very specifically to the ER membrane. This posits that information intrinsic to the AH is enough to provide membrane specificity for binding. However, we can not exclude since the ER is the largest expanse of membranes in the cell, it may be that special features are not required to bind to it, but rather certain features are required to provide specificity to other, less abundant, organelles. Regardless, our work shows that certain traits such as length and charge at the edge of the hydrophobic surface are major determinants of specificity. We hypothesize that membranes with different properties such as changes in curvature and surface charge, will attract AHs of varying lengths and aa composition, but further experimentation is required to support this.

Altogether, our data highlight an additional, previously uncharacterized, targeting motif for ER proteins. This provides the first mechanistic glimpse into the previously unexplored question of how SuPs, many of which are essential for secretory pathway functions, reach their correct destination on the ER membrane. We find factors both in trans and in cis that affect their targeting and through verifying it found conditions that reduce viral replication in its natural host cell. However, clearly much more awaits discovery. More broadly, the unique way by which SuPs specifically recognize the ER membrane highlights the complex and multifaceted nature of the processes required to ensure correct localization of the proteome and this has implications ranging from basic cell biology, through to virology and medicine.

## Figure legends

Supplementary figure 1: Human homologs of Sec31 and Dsl1 also have predicted AHs

Helical wheels of the predicted AHs in the human homologs of Sec31 (SEC31B) and Dsl1 (ZW10). Hydrophobic moment values are presented in Table S5.

## Materials and methods

### Yeast growth

Yeast cells were grown on solid media containing 2.2% agar or liquid media. YPD (2% peptone, 1% yeast extract, 2% glucose) was used for cell growth if only antibiotic selections were required, whereas synthetic minimal media (S; 0.67% [w/v] yeast nitrogen base (YNB) without amino acids and with ammonium sulphate), with 2% [w/v] glucose or galactose, supplemented with required amino acids, was used for auxotrophic selection, and for microscopy imaging. Antibiotic concentrations were as follows: nourseothricin (NAT, Jena Bioscience) at 0.2g/l and G418 (Formedium) at 0.5g/l. When G418 was added to S medium, it contained 0.17% [w/v] YNB without ammonium sulfate and with monosodium glutamate (MSG) instead to ensure basic pH suitable for G418 selection.

### Yeast strains, plasmids, and primers

All the strains used in this study are *S. cerevisiae* based on laboratory strain BY4741 (Baker Brachmann et al., 1998). Genetic transformations were performed by the lithium-acetate, polyethylene glycol, and single-stranded DNA method (Daniel Gietz and Woods, 2002).

The list of strains and plasmids used in this study can be found in Tables S6 and S7, respectively. The primers that were used for transformation in the study were designed using Primers4Yeast (Yofe and Schuldiner, 2014) and are shown in Table S8.

### Library preparation

To make the library used in the study, we used automated mating, sporulation and selection methods (described in (Cohen and Schuldiner, 2011)). The steps of this processes were done using a RoToR bench-top colony arraying instrument (Singer Instruments, UK). For the BMV 1aHB-GFP screen that is described in Figure 1, the query strain was YMS6006 (an SGA ready strain expressing BMV 1a HB-GFP) and the library was the combined knockout (KO) (Giaever et al., 2002) and DAmP (Breslow et al., 2008) collections.

### Automated high-throughput fluorescence microscopy

Automated microscopy was used to screen the deletion and DAmP library with 1aHB-GFP (Figure 1E-F). For this, the library was transferred from 1536 agar plates to 384-well polystyrene plates containing 100µl selection medium (SD(MSG) +Nat +G418 +complete AAs) and incubated overnight in a Liconic incubator at 30°C. Then, 5µl of cells were back diluted in 95µl of YPD in 384-well plates. After 4 hours, 50µl from each well was transferred to a glass-bottomed 384-well microscopy plate (Matrical Bioscience) coated with Concanavalin A (ConA). After 20 minutes of incubation at 25°C, the plates were washed two times with imaging medium (SD -riboflavin +complete AAs) and imaged using a 60X air lens (NA 0.9) and with an ORCA-flash 4.0 digital camera (Hamamatsu). Images were recorded with 488nm laser illumination for the GFP channel. All the steps in the automated imaging process were performed by an EVO freedom liquid handler (TECAN). The microscopy images were cropped, colored, and slightly adjusted for brightness/contrast using ImageJ (Schneider et al., 2012).

### Manual fluorescence microscopy

The images taken outside of the high-throughput screen (all figures except Figure 1 E, F) were done so with enhanced-resolution (SoRa) microscopy, using an Olympus IXplore SpinSR system, composed of an Olympus IX83 inverted microscope scanning unit (SCU-W1) coupled to a high-resolution spinning disk module (Yokogawa CSU-W1 SoRa confocal scanner with double microlenses and 50µm pinholes), operated by cellSens Dimension (version 3.1). The imaging medium was S +Galactose +complete AAs, to induce expression of the helices under control of the *GAL1pr*. The images were analyzed as described in the section above. Images were recorded with 488nm laser illumination for the GFP channel, and 561nm laser illumination for the mCherry channel.

### Helix prediction

The Heliquest program was used to predict AHs (Gautier et al., 2008), using the default parameters for alpha helices, with no blacklist. Structures of all of the candidates were manually screened on UniProt (UniProt Consortium, 2021), with either PDB or Alphafold structures, or both, depending on availability. Sequences with both predictions fromHeliquest and alpha helices according to structure prediction were considered the better candidates, particularly if they were annotated by Heliquest as ‘Possible lipid binding helix’ or had a prediction from an additional program. The helical wheels were created by the Heliquest tool.

### BMV 1a expression test

BMV replication was launched by using plasmids pB12VG1 and pB3VG128 (Zhang et al., 2012). Yeast cells were harvested between 0.4-1.0 OD_600_. Total protein was extracted from 2 OD_600_ units of cells as described previously (Li et al., 2016). Equal volumes of the extracted protein samples were used for SDS-PAGE and then transferred to polyvinylidene difluoride (PVDF) membrane. Target proteins were detected using a 1:10,000 dilution of rabbit anti-BMV 1a (Kind gift from Dr. Paul Ahlquist) and a 1:10,000 dilution of mouse anti-PGK1 (Thermo Fisher Scientific, #459250). Detection was performed using HRP-conjugated secondary antibodies (1:6,000 dilution, Thermo Fisher Scientific, #32430 for anti-mouse and #32460 for anti-rabbit) and Supersignal West Femto Substrate (Thermo Fisher Scientific, #34095). Protein bands were then visualized using Azure c4000 (Azure biosystems) and quantified using ImageJ.

### BMV replication test

Total RNA was extracted using the hot phenol method (Köhrer and Domdey, 1991). Equal amounts of total RNA were used for formaldehyde-agarose gel electrophoresis and then transferred onto a Nytran nylon membrane. BMV positive- and negative-strand RNAs and 18S rRNA were detected using specific ^32^P-labeled probes. Negative-strand RNA membranes were exposed longer than other Northern blot membranes.

### Conservation analysis

Putative orthologous genes of Sec31 were identified by a blast search against the uniport database which found 250 orthologous proteins, all bearing the gene name “Sec31” and the BLAST e-value was below 10^−169^. Subsequently, we employed CLUSTALX for multiple sequencing alignment followed by hierarchical clustering of the % identity matrix. Proteins clustered with *S. cerevisiae* are depicted in Figure 3A.

## Acknowledgments

We thank Yeynit Asraf, Zohar Gazi and Amir Fadel for help with screening and imaging, and Reut-Ester Avraham and Enas Katawi for technical support. We thank Rosario Valenti for her help with making the 3D models of the helices. We are thankful to Bruno Anthony, Anthony Futerman and Tamir Dingjan for fruitful discussions. We thank Dr. Paul Ahlquist for kindly sharing rabbit anti-BMV 1a antibody.

This work was supported by a collaborative grant to XW and MS (BARD US-5389-21C). Emma Fenech was supported by the Weizmann Institute of Science senior postdoctoral award. All targeting work in the Schuldiner lab is supported by an ERC CoG from the European Union (OnTarget, 864068). The robotic system of the Schuldiner lab was purchased through the kind support of the Blythe Brenden-Mann Foundation. MS is an Incumbent of the Dr. Gilbert Omenn and Martha Darling Professorial Chair in Molecular Genetics.

